# Revaluation of magnetic properties of Magneto, MagR and αGFP−TRPV1/GFP−ferritin

**DOI:** 10.1101/737254

**Authors:** Guangfu Wang, Peng Zhang, Suresh K. Mendu, Yali Wang, Yajun Zhang, Xi Kang, Bimal N. Desai, J. Julius Zhu

**Affiliations:** Department of Pharmacology, University of Virginia School of Medicine, Charlottesville, VA 22908, USA

**Author notes:** These authors contributed equally to this work (Guangfu Wang, Peng Zhang, Suresh K. Mendu). Correspondence and requests for materials should be addressed to Bimal Desai and J. Julius Zhu, Department of Pharmacology, University of Virginia School of Medicine 1300 Jefferson Park Avenue Charlottesville, VA 22908.

## Abstract

Magnetic control of neuronal activity offers many obvious advantages over electric, optogenetic and chemogenetic manipulations. A recent series of highly visible papers reported the development of magnetic actuators (i.e., Magneto, MagR and αGFP−TRPV1/GFP−ferritin) that appeared to be effective in controlling neuronal firing^1–3^, yet their action mechanisms seem to conflict with the principles of physics^4^. We found that neurons expressing Magneto, MagR and αGFP−TRPV1/GFP−ferritin did not respond to magnetic stimuli with any membrane depolarization (let alone action potential firing), although these neurons frequently generated spontaneous action potentials. Because the previous study did not establish the precise temporal correlation between magnetic stimuli and action potentials in recorded neurons^1–3^, the reported magnetically-evoked action potentials are likely to represent mismatched spontaneous firings.

## RESULTS

To examine the membrane surface incorporation of Magneto, we transfected 293T cells with the P2A-linked wild type TRPV4 (the primogenitor of Magneto2.0), ferritin (the other key element of Magneto2.0) and mCherry, aka TRPV4-P2A-ferritin-P2A-mCherry, or the P2A-linked Magneto2.0 and mCherry, aka Magneto-P2A-mCherry. We then made simultaneous measurements of the magnetic stimulation- and agonist-evoked responses in control non-expressing and TRPV4-P2A-ferritin-P2A-mCherry or Magneto-P2A-mCherry expressing cell pairs (**Fig 1a**). To determine the precise timing of applied magnetic field, we used an LED illuminator and a photodetector to monitor the exact position of magnets mounted on a Luigs-Neumann JUNIOR COMPACT manipulator. Delivering a K&J N42 neodymium 1/16” block magnet to the position 1,000 μm away from recorded cells generated a 64.5-mT magnetic field (**Fig S1**). As expected, delivery and withdrawal of the magnet did not induce any current in control and TRPV4-P2A-ferritin-P2A-mCherry expressing cells (**Fig 1b-c**). In contrast, puff application of TRPV4 agonist, GSK1016790A (**GSK101**), reliably elicited inward currents in TRPV4-P2A-ferritin-P2A-mCherry expressing cells, but not control non-expressing cells (**Fig 1b-c**). Additional analysis revealed that GSK101-elicited currents in TRPV4-P2A-ferritin-P2A-mCherry expressing cells had the IV relationship typical of TRPV4, and the currents were blocked by a TRPV4 antagonist GSK205 (**Fig S2a-b**), indicating TRPV4-specific currents^5^. Surprisingly, neither the magnetic stimuli nor GSK101 induced any significant current in control and Magneto-P2A-mCherry expressing cells (**Fig 1b-c**). Together, these results suggest that unlike wild type TRPV4, the ferritin-conjugated variant of TRPV4, i.e., Magneto2.0, fails to form a functional ion channel and/or incorporate into the plasma membrane of 293T cells.

**Figure 1.**
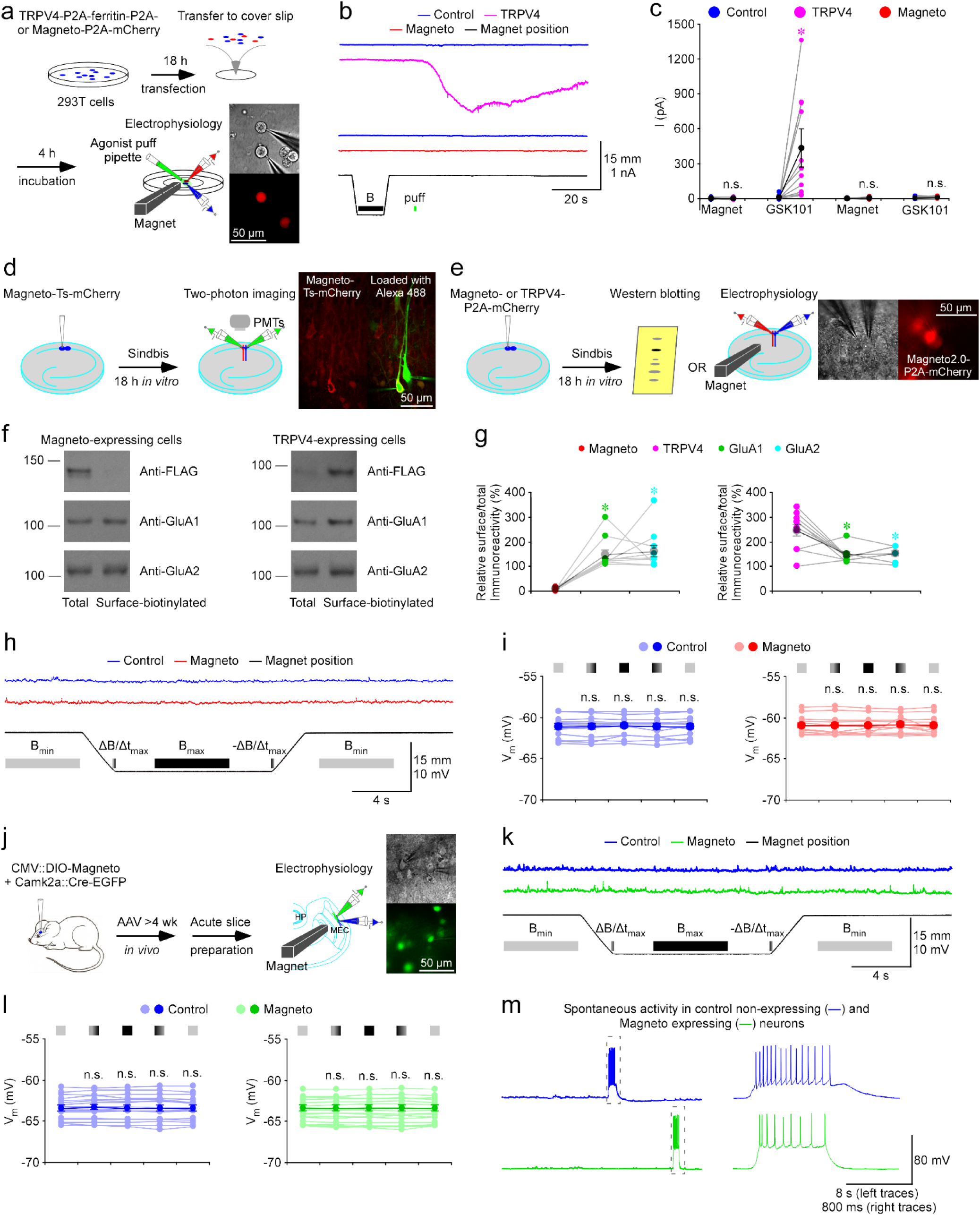
No magnetic effect in cells or neurons expressing Magneto2.0. (a) Schematic drawing outlines the design of *in vitro* transfection, magnetic stimulation and electrophysiological recordings in TRPV4-P2A-ferritin-P2A-mCherry and Magneto-P2A-mCherry expressing cultured 293T cells. The right images show simultaneous whole-cell recordings from a pair of control non-expressing and Magneto-P2A-mCherry expressing cells under transmitted light (left) and fluorescence microscopy with RFP filter (right). (b) Current recordings from neighboring control non-expressing and TRPV4-P2A-ferritin-P2A-mCherry expressing cells (left), and control non-expressing and Magneto-P2A-mCherry expressing cells (right) during magnetic stimulation and puff application of 100 nM TRPV4 agonist GSK1016790A (GSK101). (c) Values of currents of control non-expressing and TRPV4-P2A-ferritin-P2A-mCherry expressing 293T cells during magnetic stimulation and puff application of GSK101 (Ctrl: −0.5±0.9 pA; EXP: −1.8±1.1 pA, *Z*=-1.511, *p*=0.12 for control cells; Ctrl: 9.0±5.3 pA; EXP: 433.5±132.6 pA, *Z*=2.934, *p*<0.005 for TRPV4 expressing cells; *n*=11). Values of currents of control non-expressing and Magneto-P2A-mCherry expressing 293T cells during magnetic stimulation and puff application of GSK101 (Ctrl: −0.2±0.6 pA; EXP: 0.6±1.4 pA, *Z*=-0.175, *p*=0.86 for control cells; Ctrl: 4.1±2.2 pA; EXP: 7.1±2.0 pA, *Z*=1.293, *p*=0.20 for Magneto expressing cells; *n*=13). Asterisk indicates *p*<0.05 (Wilcoxon tests). (d) Schematic drawing outlines the design of *in vitro* Sindbis viral expression and two-photon imaging in cultured rat hippocampal slices. The right two-photon images show a pair of control non-expressing and Magneto-Ts-mCherry expressing CA1 pyramidal neurons after loading Alexa 488 with patch-clamp pipettes (left: mCherry channel only; right: mCherry and Alexa 488 red channels overlay). (e) Schematic drawing outlines the design of *in vitro* Sindbis viral expression, biochemistry analysis, magnetic stimulation and electrophysiological recordings in cultured rat hippocampal slices. The right images show simultaneous whole-cell recordings from a pair of control non-expressing and Magneto-P2A-mCherry expressing CA1 pyramidal neurons under transmitted light (left) and fluorescence microscopy with RFP filter (right). (f) Western blots of total and membrane surface-biotinylated recombinant Magneto2.0 and TRPV4 (both of which are FLAG tagged), and endogenous GluA1 and GluA2 in CA1 cells prepared from cultured rat hippocampal slices. Each lane loaded with 20 μg proteins. (g) Relative levels of membrane surface-biotinylated vs. total Magneto2.0 (Magneto2.0: 7.7±1.3%; GluA1: 148.0±17.8%, *n*=11, *Z*=2.934, *p*<0.005; GluA2: 160.2±23.3%; *n*=11, *Z*=2.934, *p*<0.005) and TRPV4 (TRPV4: 245.4±24.0%; GluA1: 145.5±9.4%, *n*=10, *Z*=-2.396, *p*<0.05; GluA2: 152.7±10.1%, *n*=10, *Z*=-2.396, *p*<0.05) compared to GluA1 and GluA2. Asterisks indicate *p*<0.05 (Wilcoxon tests). (h) Recordings of membrane potentials of the pair of control non-expressing and Magneto-P2A-mCherry expressing CA1 pyramidal neurons before, during and after magnetic stimuli delivered with a K&J N42 1/16” permanent block magnet mounted on a micromanipulator. (i) Values of membrane potentials of control non-expressing (Initial B_min_: −61.1±0.3 mV; ΔB/Δt_max_: −61.2±0.4 mV, *Z*=-1.038, *p*=0.28; B_max_: −61.1±0.3 mV, *Z*=-0.105, *p*=0.92; -ΔB/Δt_max_: −61.2±0.4 mV, *Z*=-0.364, *p*=0.70; Ending B_min_: −61.1±0.3 mV, *Z*=1.083, *p*=0.28; Wilcoxon tests) and Magneto-P2A-mCherry (Initial B_min_: −60.9±0.3 mV; ΔB/Δt_max_: −61.0±0.3 mV, *Z*=-0.664, *p*=0.51; B_max_: −61.0±0.3 mV, *Z*=-1.103, *p*=0.31; -ΔB/Δt_max_: −60.8±0.3 mV, *Z*=-0.314, *p*=0.75; Ending B_min_: −61.0±0.3 mV, *Z*=-0.734, *p*=0.46; Wilcoxon tests) expressing CA1 pyramidal neurons when the permanent magnet was away from (light), approaching to (light-dark transient color), close to (dark), retracting from (dark-light transient color), and away from (light) recorded neurons (*n*=13 from 6 animals). Note dark lines/dotes indicating averages and shaded lines/dotes indicating experimental data. Note also no difference in membrane potential in control non-expressing and Magneto-P2A-mCherry expressing CA1 pyramidal neurons in all the experimental stages (*p*>0.05; Wilcoxon tests). (j) Schematic drawing outlines the design of *in vivo* AAV viral expression of CMV::DIO-Magneto and Camk2a::Cre-EGFP, *ex vivo* magnetic stimulation and electrophysiological recordings in acutely prepared mouse MEC slices. The right images show simultaneous whole-cell recordings from a pair of control non-expressing and DIO-Magneto/Cre-GFP expressing MEC layer 2/3 pyramidal neurons under transmitted light (left) and fluorescence microscopy with GFP middle filter (right). (k) Recordings of membrane potentials of the pair of control non-expressing and DIO-Magneto/Cre-GFP expressing MEC layer 2/3 neurons before, during and after magnetic stimuli delivered with a K&J N42 1/16” permanent block magnet mounted on a micromanipulator. (l) Values of membrane potentials of control non-expressing (Initial B_min_: −63.4±0.3 mV; ΔB/Δt_max_: −63.4±0.4 mV, *Z*=1.018, *p*=0.31; B_max_: −63.4±0.4 mV, *Z*=0.213, *p*=0.83; -ΔB/Δt_max_: −63.4±0.4 mV, *Z*=0.734, *p*=0.46; Ending B_min_: −63.4±0.3 mV, *Z*=-1.065, *p*=0.29) and Cre-GFP/DIO-Magneto2.0 (Initial B_min_: −63.4±0.3 mV; ΔB/Δt_max_: −63.4±0.4 mV, *Z*=-0.024, *p*=0.98; B_max_: −63.4±0.4 mV, *Z*=0.166, *p*=0.87; -ΔB/Δt_max_: −63.4±0.4 mV, *Z*=0.166, *p*=0.87; Ending B_min_: −63.4±0.3 mV, *Z*=-1.207, *p*=0.23) expressing MEC L2/3 pyramidal and stellate neurons when the permanent magnet was away from (light), approaching to (light-dark transient color), close to (dark), retracting from (dark-light transient color), and away from (light) recorded neurons (*n*=17 from 11 animals). Note dark lines/dotes indicating averages and shaded lines/dotes indicating experimental data. Note also no difference in membrane potential in control non-expressing and DIO-Magneto/Cre-GFP expressing MEC L2/3 neurons in all the experimental stages (*p*>0.05; Wilcoxon tests). (m) Recordings of spontaneous events in the pair of control non-expressing and DIO-Magneto/Cre-GFP expressing MEC layer 2/3 neurons. Note that the spontaneous suprathreshold events in the gray dash line boxes are shown again in an expanded time scale in the right.

To test whether Magneto might evoke action potentials in neuronal cells, we first expressed mCherry-fused Magneto2.0, aka Magneto-Ts-mCherry, and Magneto-P2A-mCherry in CA1 neurons of cultured rat hippocampal slices using the established Sindbis viral expression system (**Fig 1d**; see^6, 7^ for the methods). Two-photon images showed that Magneto-Ts-mCherry, although robustly expressed, seemed to have limited, if any, presence at the plasma membrane of CA1 neurons (**Fig 1d insets**). In consistent, Western blots showed that in contrast to TPRV4 in TRPV4-P2A-ferritin-P2A-mCherry expressing CA1 cells, Magneto2.0 membrane surface expression was minimal despite its high intracellular expression in Magneto-P2A-mCherry expressing CA1 cells (**Fig 1e-g**). To verify the results, we expressed Magneto-P2A-mCherry and TRPV4-P2A-ferritin-P2A-mCherry in the mouse barrel cortex *in vivo* for 7-10-days using the established lentiviral expression system (**Fig S3**; see^6^ for the methods). Again, in spite of high intracellular expressions, only TPRV4, but not Magneto2.0, showed efficient membrane surface expression in barrel cortical neurons (**Fig S3**). These results are consistent with the fact that C-terminus of the Magneto2.0 primogenitor TRPV4, which is essential for surface trafficking and functional expression of TRPV4^5^, is deleted in Magneto2.0^2^. We then made simultaneous whole-cell recordings from pairs of control non-expressing and Magneto-P2A-mCherry expressing CA1 cells. Patch-clamp recordings showed that application of up to 64.5 mT static magnetic field induced neither depolarization nor action potential firing in control and Magneto-P2A-mCherry expressing CA1 cells (**Fig 1h-i**). Similarly, the magnetic stimuli failed to induce depolarization and action potential discharge in Magneto-P2A-mCherry expressing CA1 cells when the expression was made with lentivirus (**Fig S4**). These results indicate that the magnetic stimuli do not induce action potential in Magneto expressing CA1 neurons in cultured slices.

To further examine Magneto, we made *in vivo* Sindbis viral expression of Magneto-P2A-mCherry in layer 2/3 pyramidal and stellate neurons in the mouse medial entorhinal cortex (**MEC**) or layer 5 pyramidal neurons in the mouse barrel cortex for ∼18 hrs, and then acutely prepared entorhinal or barrel cortical slices (**Figs S5a and S6a**). Simultaneous recordings showed that the 64.5 mT static magnetic field did not induce any depolarization or action potential firing in control and Magneto-P2A-mCherry expressing neurons (**Figs S4 and S6**). We repeated the experiments using the same 3/8” permanent block magnet employed in the previous study^2^ (**Fig S7a**). Positioning the magnet 5.00 mm away from recorded neurons, which yielded a 78.8 mT static magnetic field (**Fig S1**), induced neither depolarization nor action potential firing in control and Magneto-P2A-mCherry expressing entorhinal neurons (**Fig S7b-c**). In these experiments, we noted that control and Magneto-P2A-mCherry expressing entorhinal neurons frequently displayed spontaneous synaptic events, and at times, the spontaneous events reached the threshold and triggered bursts of action potentials (**Figs S5b, S5d and S7b**), suggest a potential cause for the reported magnetic effects^2^.

We then recorded hippocampal neurons after ∼3‒5-week *in vivo* AAV viral expression of DIO-Magneto and GFP-Cre (**Fig S8a**), using the same brain slice tissues prepared and published in *Wheeler et al.*^2^. The 78.8 mT static magnetic field induced neither depolarization nor action potential discharge in control or Magneto expressing dentate gyrus neurons (**Fig S8b-c**). Finally, we made *in vivo* AAV viral expression of DIO-Magneto and GFP-Cre in layer 2/3 pyramidal and stellate neurons of mouse MEC for ∼3-5 weeks, and then recorded activity of entorhinal neurons in acutely prepared entorhinal cortical slices (**Fig 1j**). Once again, simultaneous recordings showed that application of up to 64.5 mT static magnetic field did not induce any depolarization or action potential firing in control and DIO-Magneto/GFP-Cre expressing entorhinal neurons (**Fig 1k-l**). Importantly, we observed abundant spontaneous activities that from time to time, reached the firing threshold and elicited action potentials in these experiments (**Figs 1k, 1m, S8b and S8d**). Collectively, our results consistently support the idea that Magneto does not function as an effective magnetic actuator and spontaneous action potentials can confound the interpretation of Magneto expressing neurons subjected to magnetic stimuli.

The above experiments raised the concern about the other two recently reported magnetic actuators, MagR and αGFP−TRPV1/GFP−ferritin, given no time correlation between magnetic stimuli and action potentials in MagR and αGFP−TRPV1/GFP−ferritin expressing neurons reported in the two other studies^1, 3^. To test whether MagR may serve as a magnetic actuator, we made Sindbis viral expression of MagR-P2A-GFP in CA1 neurons of cultured rat hippocampal slices (**Fig 2a**). After 18-hour expression, we made simultaneous whole-cell recordings from pairs of control non-expressing and MagR-P2A-GFP expressing CA1 cells. Application of up to 64.5 mT static magnetic field induced neither depolarization nor action potential firing in control and MagR-P2A-GFP expressing CA1 cells (**Fig 2b-c**). These results are indicative of no magnetic effect on MagR expressing CA1 neurons, and the previously reported MagR-mediated action potentials to be confounding spontaneous firing (cf.^8^).

**Figure 2.**
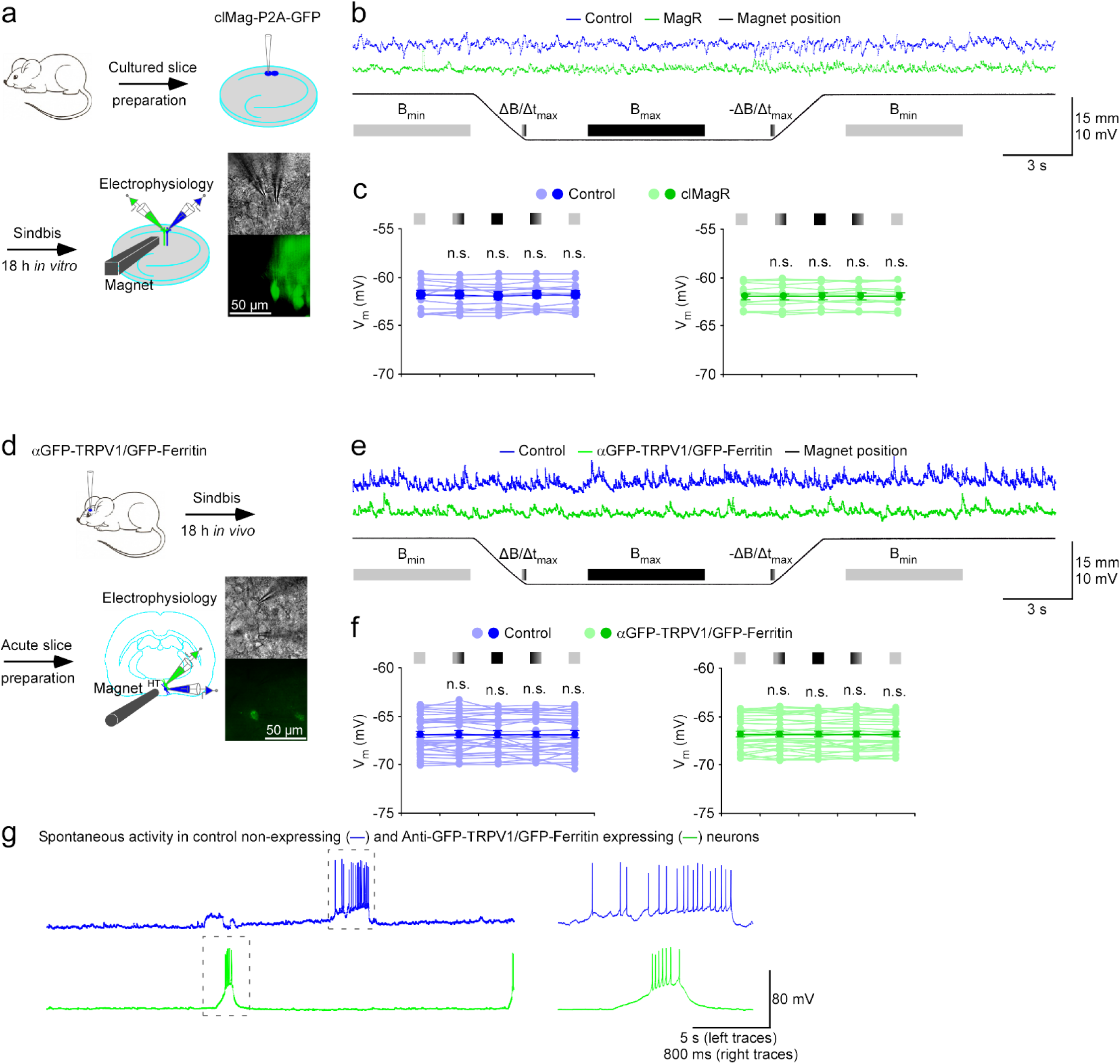
No magnetic effect in neurons expressing MagR or αGFP-TRPV1/GFP-ferritin. (a) Schematic drawing outlines the design of *in vitro* Sindbis viral expression, magnetic stimulation and electrophysiological recordings in cultured rat hippocampal slices. The right images show simultaneous whole-cell recordings from a pair of control non-expressing and MagR expressing CA1 pyramidal neurons under transmitted light (left) and fluorescence microscopy with GFP middle filter (right). (b) Recordings of membrane potentials of the pair of control non-expressing and MagR expressing CA1 pyramidal neurons before, during and after magnetic stimuli delivered with a K&J N42 1/16” permanent block magnet mounted on a micromanipulator. (c) Values of membrane potentials of control non-expressing (Initial B_min_: −62.0±0.4 mV; ΔB/Δt_max_: −62.0±0.4 mV, *Z*=-0.934, *p*=0.35; B_max_: −62.0±0.4 mV, *Z*=-0.804, *p*=0.42; -ΔB/Δt_max_: −61.9±0.4 mV, *Z*=0.804, *p*=0.42; Ending B_min_: −62.0±0.4 mV, *Z*=0.035, *p*=0.97; Wilcoxon tests) and MagR (Initial B_min_: −62.0±0.3 mV; ΔB/Δt_max_: −62.0±0.3 mV, *Z*=0.175, *p*=0.86; B_max_: −61.9±0.3 mV, *Z*=1.433, *p*=0.15; -ΔB/Δt_max_: −61.9±0.3 mV, *Z*=1.013, *p*=0.31; Ending B_min_: −62.0±0.3 mV, *Z*=0.874, *p*=0.38; Wilcoxon tests) expressing CA1 pyramidal neurons when the permanent magnet was away from (light), approaching to (light-dark transient color), close to (dark), retracting from (dark-light transient color), and away from (light) recorded neurons (*n*=13 from 6 animals). Note dark lines/dotes indicating averages and shaded lines/dotes indicating experimental data. Note also no difference in membrane potential in control non-expressing and MagR expressing CA1 pyramidal neurons in all the experimental stages (*p*>0.05; Wilcoxon tests). (d) Schematic drawing outlines the design of *in vivo* Sindbis viral expression, *ex vivo* magnetic stimulation and electrophysiological recordings in acutely prepared mouse hypothalamic slices. The right images show simultaneous whole-cell recordings from a pair of control non-expressing and αGFP-TRPV1/GFP-ferritin expressing hypothalamic neurons under transmitted light (left) and fluorescence microscopy with GFP middle filter (right). (e) Recordings of membrane potentials of the pair of control non-expressing and αGFP-TRPV1/GFP-ferritin expressing hypothalamic neurons before, during and after magnetic stimuli delivered with a K&J N52 1/16” permanent cylinder magnet mounted on a micromanipulator. (f) Values of membrane potentials of control non-expressing (Initial B_min_: −67.0±0.3 mV; ΔB/Δt_max_: −66.9±0.3 mV, *Z*=0.999, *p*=0.32; B_max_: −66.9±0.3 mV, *Z*=0.020, *p*=0.98; -ΔB/Δt_max_: −67.0±0.4 mV, *Z*=0.078, *p*=0.94; Ending B_min_: −67.0±0.3 mV, *Z*=-0.549, *p*=0.58) and αGFP-TRPV1/GFP-ferritin (Initial B_min_: −66.9±0.3 mV; ΔB/Δt_max_: −66.8±0.3 mV, *Z*=0.588, *p*=0.56; B_max_: −66.9±0.3 mV, *Z*=0.020, *p*=0.98; -ΔB/Δt_max_: −66.9±0.3 mV, *Z*=0.353, *p*=0.72; Ending B_min_: −66.9±0.3 mV, *Z*=-0.882, *p*=0.38) expressing hypothalamic neurons when the permanent magnet was away from (light), approaching to (light-dark transient color), close to (dark), retracting from (dark-light transient color), and away from (light) recorded neurons (*n*=31 from 13 animals). Note dark lines/dotes indicating averages and shaded lines/dotes indicating experimental data. Note also no difference in membrane potential in control non-expressing and αGFP-TRPV1/GFP-ferritin expressing hypothalamic neurons in all the experimental stages (*p*>0.05; Wilcoxon tests). (g) Recordings of spontaneous events in the pair of control non-expressing and αGFP-TRPV1/GFP-ferritin expressing hypothalamic neurons. Note that the spontaneous suprathreshold events in the gray dash line boxes are shown again in an expanded time scale in the right.

We next examined whether αGFP−TRPV1/GFP−ferritin may serve as a magnetic actuator. In this experiment, we made Sindbis viral expression of αGFP−TRPV1-P2A-GFP−ferritinin hypothalamic neurons in intact mouse brains for 18 hours, and then made simultaneous whole-cell recordings from pairs of control non-expressing and αGFP−TRPV1/GFP−ferritin expressing hypothalamic neurons in acutely prepared slices (**Fig 2d**). Following the previous report^3^, we delivered a K&J N52 neodymium 1/16” cylinder magnet to the position 500 μm away from recorded cells, which generated a 108.0-mT magnetic field (**Fig S1**). Application of up to 108.0 mT static magnetic induced neither depolarization nor action potential discharge in control and αGFP−TRPV1/GFP−ferritin expressing hypothalamic neurons (**Fig 2e-f**). Both control and αGFP−TRPV1/GFP−ferritin expressing hypothalamic neurons exhibited frequent spontaneous synaptic potentials, and at times, depolarizing spontaneous potentials reached the threshold to trigger a burst of action potentials (**Figs 2e and 2g**). These results are suggestive of no magnetic effect on αGFP−TRPV1/GFP−ferritin expressing hypothalamic neurons and the previously reported αGFP−TRPV1/GFP−ferritin-mediated responses to be false positive magnetic effects caused by spontaneous activity.

## DISCUSSION

In summary, we systematically investigated whether Magneto2.0 might function as a magnetic actuator with multiple approaches (i.e., transfection, Sindibis, lentivirus, and AAV viral expression *in vitro* and/or *in vivo*), multiple cell types (i.e., 293T cells, hippocampal CA1 neurons, dentate gyrus neurons, cortical L5 pyramidal neurons, entorhinal layer 2/3 stellate and pyramidal neurons), and multiple animal species (i.e., rats and mice). Our results, together with two accompanied studies that used additional approaches (e.g. viral expression), cell types (e.g., cerebral Purkinje neurons and barrel cortical layer 2/3 neurons), and manipulation/recording SFV methods (e.g., electric magnetic stimuli and *in vivo* recordings)^9, 10^, consistently support the notion that Magneto2.0 does not serve as a magnetic actuator. Our experiments raise serious concerns of about the experimental design and execution of *Wheeler et al.* study. For example, deleting C-terminus of the Magneto2.0 primogenitor TRPV4^2^, that is essential for surface trafficking of Magneto2.0^5^, is poorly conceived. Moreover, whether the ferritin subunit-fused TRPV4s may multiplex together to form functional ferritins remains to be tested. Notably, *Wheeler et al.* used the recorded action potentials as the primary evidence to argue that Magneto2.0 serves as a magnetic actuator (and its surface expression)^2^, yet these electrophysiology recordings were poorly controlled and confounded by spontaneous action potentials. Thus, the rigorous electrophysiology experiment that is essential for validating Magneto as a magnetic actuator remains missing (also see ^11^ with data suggestive of the endoplasmic reticulum as the source of minimal intracellular Ca^2+^ elevation). Indeed, with the location of stimulating magnets continuously monitored, multiple independent electrophysiological experiments in this study (some using the exact same magnetic stimulations and/or tissues prepared for *Wheeler et al.*), together with those in the two accompanied studies^9, 10^, consistently demonstrated that Magneto did not respond to magnetic stimuli with any membrane depolarization (let alone action potential firing). We also show here that MagR and αGFP−TRPV1/GFP−ferritin do not produce any detectable magnetically-elicited membrane depolarization. Together, these results support the theoretical conclusion that Magneto, MagR and αGFP−TRPV1/GFP−ferritin are incapable of controlling neuronal activity by producing magnetically-evoked action potentials. We hope that our comprehensive testing of Magneto establishes a sample set of criteria to aid continuing tool-engineering efforts, including building of a magnetogenetic toolbox. The criteria include: first, surface expression validation; second, functional validation; and third, electrophysiological validation. Obviously, beyond the proof-of-principle, the applicability of novel tools is best ensured by using them to answer fundamental biology questions in a definitive manner, as was typical of previous developments in patch-clamp and imaging technology.

## ACKNOWLEDGEMENTS

We thank members of the Zhu laboratory for comments and technical assistance, many colleagues around the world for their encouragement, discussion, suggestions and/or sharing unpublished Magneto data. We also thank Dr. Can Xie for supplying clMagR, Drs. Sarah Stanley and Jeffrey Friedman for supplying αGFP−TRPV1/GFP−ferritin, and Drs Chris Deppmann, Ali Güler and/or Manoj Patel for supplying TRPV4-P2A-ferritin-P2A-mCherry, Magneto-P2A-mCherry and Magneto-Ts-mCherry constructs and AAV DIO-Magneto and GFP-Cre viruses, and for the opportunity to visit and observe their electrophysiology experiments and multiple data cross-examination meetings in late 2015 after our many futile attempts to reproduce their findings. This study is supported in part by the Epilepsy Foundation postdoctoral fellowship No. 310443 (G.W.), Alzheimer’s Association research fellowship (P.Z.), NIH grants GM108989 (B.N.D.) and NS104670 (J.J.Z.). J.J.Z. is the Radboud Professor and Sir Yue-Kong Pao Chair Professor.

## AUTHOR CONTRIBUTIONS

G.W., P.Z. and S.K.M. carried out the experiments with assistance from Y.W, Y.Z. and X.K.; B.N.D. and J.J.Z. wrote the manuscript with input from all coauthors.

## METHODS

### Animal preparation

Male and female Sprague Dawley rats and C57BL/6 mice were used to prepare cultured slices and acute slices used in this study. Animals were maintained in the animal facility at the University of Virginia and family or pair housed in the temperature-controlled animal room with 12-h/12-h light/dark cycle. Food and water were available *ad libitum*. All procedures for animal surgery and maintenance were performed following protocols approved by the Animal Care & Use Committee of the University of Virginia and in accordance with US National Institutes of Health guidelines.

### Cultured slice preparation

Cultured slices were prepared from postnatal 6−7 day old rats or mice (P6−7) as reported in our previous studies^6, 7^. In brief, the hippocampi were dissected out in ice-cold HEPES-buffered Hanks’ solution (pH 7.35) under sterile conditions, sectioned into 400-µm slices on a tissue chopper, and explanted onto a Millicell-CM membrane (0.4-µm pore size; Millipore). The membranes were then placed in 750 µl of MEM culture medium, contained (in mM): HEPES 30, heat-inactivated horse serum 20%, glutamine 1.4, D-glucose 16.25, NaHCO_3_ 5, CaCl_2_ 1, MgSO_4_ 2, insulin 1 mg/ml, ascorbic acid 0.012% at pH 7.28 and osmolarity 320. Cultured slices were maintained at 35°C, in a humidified incubator (ambient air enriched with 5% CO_2_).

### Constructs of recombinant proteins and expression

All constructs, including TRPV4-P2A-ferritin-P2A-mCherry, Magneto-P2A-mCherry and Magneto-Ts-mCherry were generously supplied by Drs Chris Deppmann and Ali Güler. Magneto-P2A-mCherry and Magneto-Ts-mCherry were subcloned into Sindbis and lenitiviral vectors. AAV viral solutions of the Cre-dependent Magneto2.0 AAV virus, aka AAV1-CMV::DIO-Magneto, and AAV9-Camk2a::EGFP-Cre were also supplied by Drs Chris Deppmann and Ali Güler. A P2A sequence was used to link clMagR^12^ and GFP, and anti-GFP−TRPV1 and GFP−ferritin^13^ to create clMagR-P2A-GFP and αGFP−TRPV1-P2A-GFP−ferritin, which were then subcloned into Sindbis viral vector. Construct expression followed our previous studies^14, 15^. For expression in cultured 293T cells, Magneto-P2A-mCherry and TRPV4-P2A-ferritin-P2A-mCherry were transfected using the calcium phosphate transfection method. For expression in cultured slices, CA1 pyramidal neurons in hippocampal cultured slices were infected after 8−18 days *in vitro* with lentivirus or Sindbis virus, and then incubated on culture media and 5% CO_2_ before experiments. For expression in intact brains, P18−28 mice were initially anesthetized by an intraperitoneal injection of ketamine and xylazine (10 and 2 mg/kg, respectively). Animals were then placed in a stereotaxic frame and one or multiple small (∼1×1 mm) holes were opened above the cortex. A glass pipette was used to make pressure injections of ∼100 nl Sindbis or lentiviral solution, or 200 nl equivolume mixture of AAV viral solutions of AAV1-CMV::DIO-Magneto and AAV9-Camk2a::EGFP-Cre into the barrel cortex, hippocampus and/or MEC according to their stereotaxic coordinates. After injection, animals were allowed to recover from the anesthesia and returned to their cages. Experiments were typically performed within 18±2 hours after Sindbis viral infection, 7−10 days after lentiviral infection and 3−5 weeks after AAV viral infection.

### Biochemical analysis

Hippocampal extracts were prepared by homogenizing hippocampal CA1 regions isolated from cultured slices, while cortical extracts were prepared by homogenizing mCherry expressing barrel cortical areas isolated from acute cortical slices. Membranes were blotted with anti-FLAG antibody (1:5,000 for *in vitro* expression, 1:2,000 for *in vivo* expression; Fisher Scientific, Hampton, NH; Cat# MA1-91878, RRID:AB_1957945), stripped and reblotted twice with anti-GluA1 (1:1,000 for *in vitro* expression; 1:1,000 for *in vivo* expression; EMD Millipore, Burlington, MA; Cat# AB1504, RRID:AB_2113602) or anti-GluA2 antibody (1:6,000 for *in vitro* expression; 1:6,000 for *in vivo* expression; EMD Millipore; Cat# AB1768, RRID:AB_2313802). Western blots were quantified by chemiluminescence and densitometric scanning of the films under linear exposure conditions.

### Electrophysiology and two-photon imaging

Simultaneous multiple whole-cell recordings were obtained from nearby expressing and non-expressing 293T cells, CA1 pyramidal neurons, barrel cortical layer 5 pyramidal neurons, dentate gyrus neurons, entorhinal layer 2/3 stellate and pyramidal neurons, under visual guidance with fluorescence and transmitted light illumination^7, 16^, using up to two Axopatch-200B (voltage clamp) or Axoclamp 2B (current clamp) amplifiers (Molecular Devices, LLC, Sunnyvale, CA). Bath solution (29±1.5°C), unless otherwise stated, contained (in mM): NaCl 119, KCl 2.5, CaCl_2_ 2, MgCl_2_ 1, NaHCO_3_ 26, NaH_2_PO_4_ 1, glucose 25, at pH 7.4 and gassed with 5% CO_2_/95% O_2_. Patch recording pipettes (3-6 MΩ) for current (voltage-clamp) recordings contained (in mM): cesium methanesulfonate 115, CsCl 20, HEPES 10, MgCl_2_ 2.5, Na_2_ATP 4, Na_3_GTP 0.4, sodium phosphocreatine 10, EGTA 0.6, and spermine 0.1, at pH 7.25; and for voltage (current-clamp) recordings contained: potassium gluconate 115, HEPES 10, MgCl_2_ 2, MgATP 2, Na_2_ATP 2, Na_3_GTP 0.3 and KCl 20, at pH 7.25.

Two-photon imaging and electrophysiology were simultaneously performed using a custom-built microscope operated by a custom-written IGOR Pro 6 program (WaveMetrics, Lake Oswego, OR)^6, 16^. Neighboring expressing and non-expressing CA1 or dentate gyrus neuron pairs were broken in simultaneously to load green indicator Alexa 488 (20 μM). Images were taken ∼15−30 minutes after loading of the indicator. Alexa 488 and mCherry were excited by a femtosecond Ti:Sapphire laser (Chameleon Ultra; Coherent, Santa Clara, CA) at a wavelength of 880 nm.

### Agonist application and magnetic stimulation

TRPV4 agonist, GSK1016790A (EMD Millipore, Billerica, MA), was puff applied with a brief (1 sec) air pressure by a glass pipette mounted on a Luigs-Neumann JUNIOR COMPACT manipulator (Luigs-Neumann GmbH, Ratingen, Germany) and positioned ∼150 µm away from the recorded 293T cells. TRPV4-specific antagonist GSK205 (EMD Millipore), was bath applied. Magnetic stimuli were made using axially magnetized magnets, including a 3/8” cylinder magnet used in the previous study^2^ and N42 1/16” x 1/4” block neodymium magnets purchased from K&J Magnetics. The magnetic intensity-distance relationships of these magnets were calculated with an HT20 Gauss Tesla meter (Shanghai Hengtong Cidian Technology Co., Ltd., Shanghai, China) (**Fig S1**). A Luigs-Neumann JUNIOR COMPACT manipulator was used to mount and rapidly position the magnets to 1.00 mm (for N42 block magnets), 0.50 mm (for N52 cylinder magnets) or 5.00 mm (for the 3/8” cylinder magnet) away from recorded cells to create a static magnetic field >50 mT or >100 mT. After stimulation, the magnets were rapidly withdrawn by 12−15 mm to eliminate magnetic stimuli. To determine the precise timing of applied magnetic field, we used an LED illuminator and a photodetector (Thorlabs Inc, Newton, NJ) to monitor the exact position of magnets mounted on a Luigs-Neumann JUNIOR COMPACT manipulator.

### Statistical analysis

Statistical results were reported as mean±s.e.m. Animals or cells were randomly assigned into control or experimental groups and investigators were blinded to experiment treatments. Given the negative correlation between the variation and square root of sample number, *n,* the group sample size was typically set to be ∼10−25 to optimize the efficiency and power of statistical tests. Statistical significances of the means (*p*<0.05; two sides) were determined using Wilcoxon non-parametric tests for paired samples. The data that support the findings of this study are available from the corresponding authors upon request.

---

**Figure S1.**
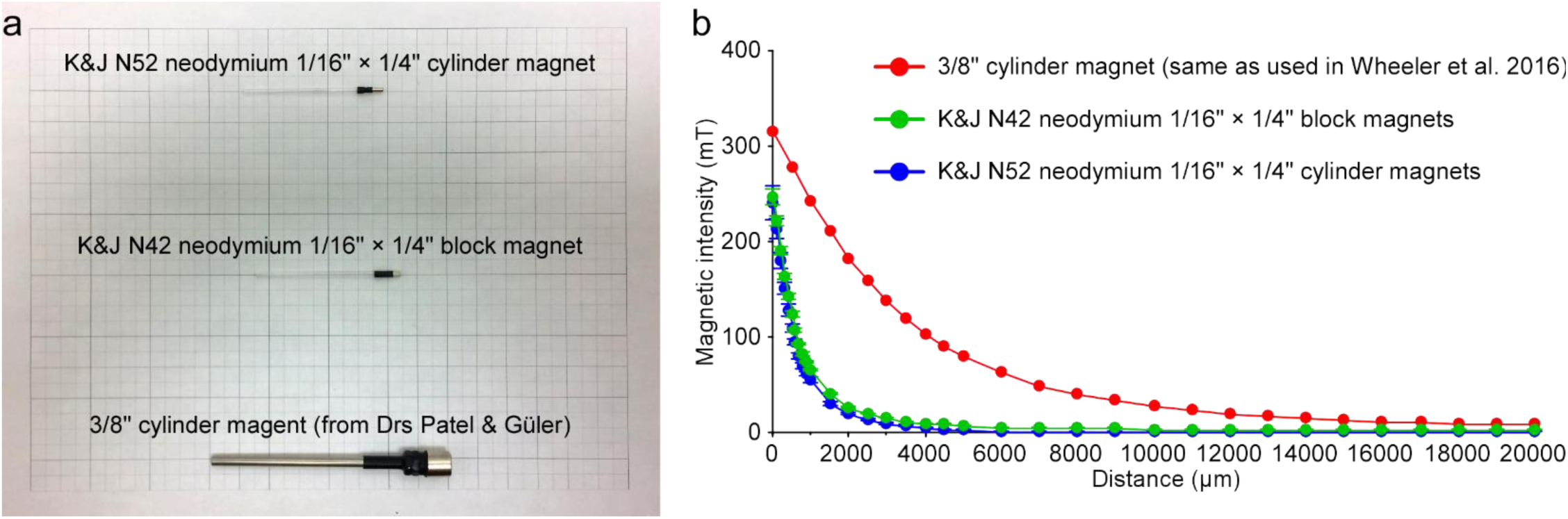
Properties of magnets used in experiments. (a) Image shows the two frequently used magnets attached to a 2 mm diameter borosilicate glass or a 4 mm diameter stainless steel rod. (b) Plots of the magnetic intensity against the distance of N42 neodymium 1/16” × 1/4” block magnets (K&J Magentics, Inc.; *n*=4) and a 3/8” cylinder magnet (the same as used in the previous report^2^; *n*=1).

**Figure S2.**
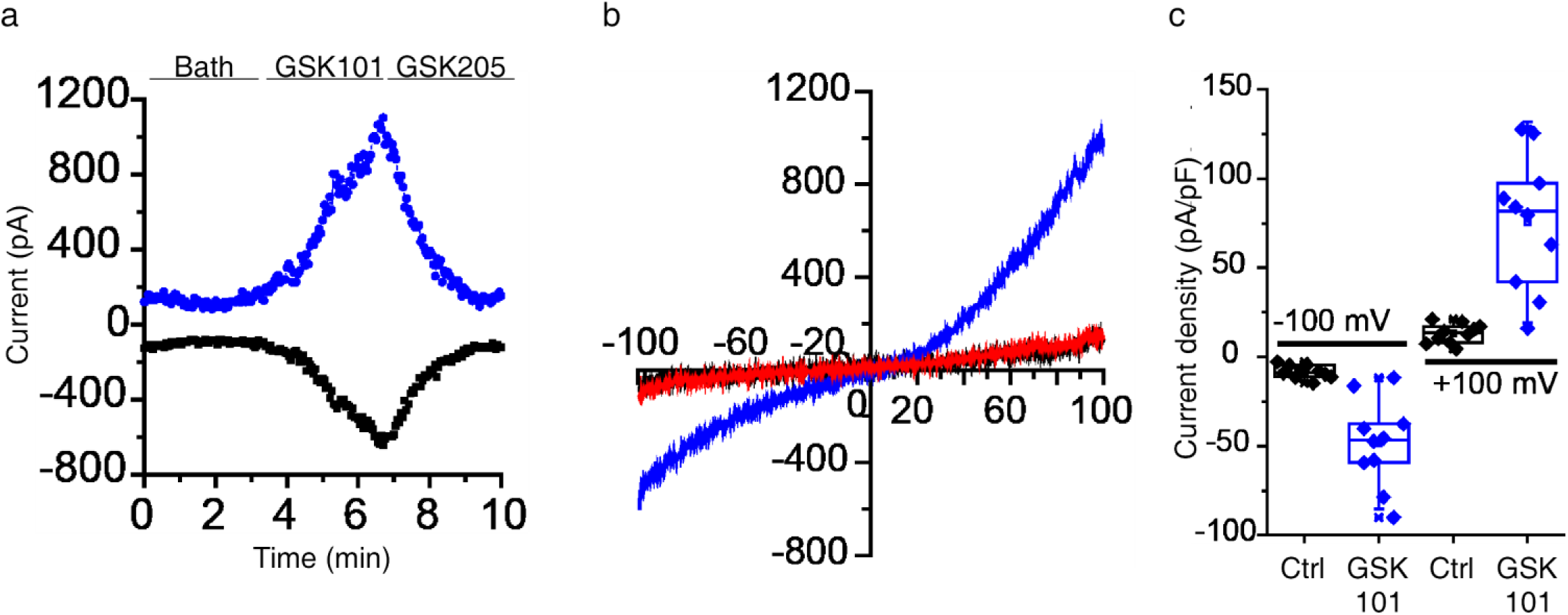
TRPV4-specific currents in TRPV4-P2A-ferritin-P2A-mCherry expressing 293T cells. (a) Whole-cell patch-clamp recordings of outward (at 100 mV, blue trace) and inward (at −100mV, black trace) currents in 293T cells expressing control TRPV4-P2A-ferritin-P2A-mCherry in response to TRPV4 agonist GSK101 (10 nM) and antagonist GSK205 (10 µM). (b) Current-voltage relationships of control (black trace), the GSK101- (blue trace) and GSK205- (red trace) evoked responses in 293T cells expressing TRPV4-P2A-ferritin-P2A-mCherry. (c) Box charts showing quantification of the peak currents in 293T cells expressing TRPV4-P2A-ferritin-P2A-mCherry (Ctrl: −8.3±1.2 pA; GSK101: −48.5±7.8 pA; *n*=10, *Z*=-2.803, *p*<0.01, for inward currents at −100 mV; Ctrl: 13.1±1.7 pA; GSK101: 75.5±11.9 pA; *n*=10, *Z*=2.803, *p*<0.01, for outward currents at 100 mV).

**Figure S3.**
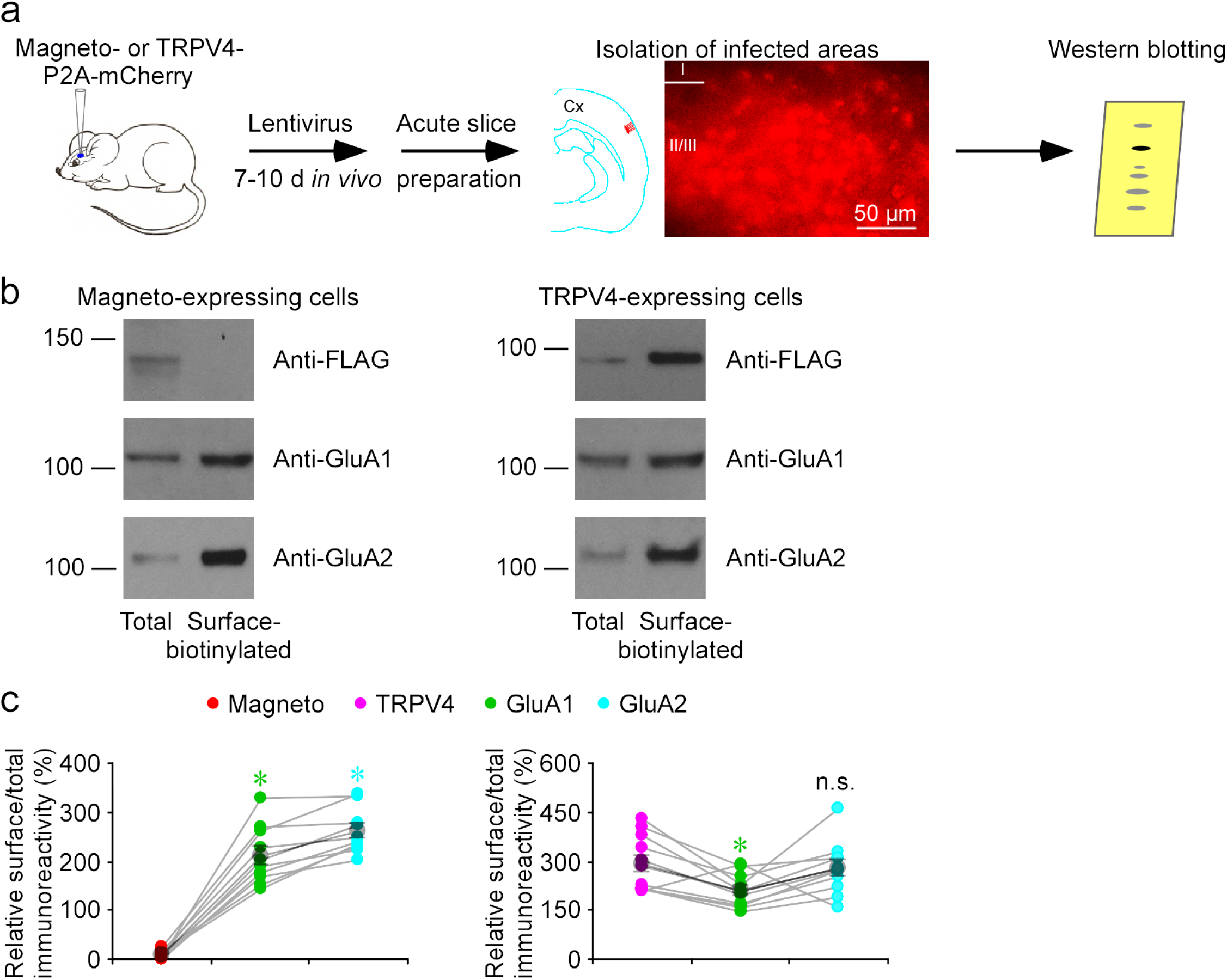
Membrane surface trafficking of Magneto2.0 is impaired in barrel cortical cells *in vivo*. (a) Schematic drawing outlines the design of biochemistry analysis of Magneto2.0 and TRPV4 (both of which are FLAG tagged) expressed in the intact mouse barrel cortex with lentivirus. The inset image shows the expressing barrel cortical area identifiable by mCherry fluorescence. Note the majority of barrel cortical neurons nearby the viral injected site expressing mCherry. (b) Western blots of total and membrane surface-biotinylated recombinant Magneto2.0 and TRPV4, and endogenous GluA1 and GluA2 in barrel cortical neurons after 7-10 days of expression. Each lane loaded with 40 μg proteins. (c) Relative levels of membrane surface-biotinylated vs. total Magneto2.0 (Magneto2.0: 9.9±2.5%; GluA1: 212.2±18.7%, *n*=10, *Z*=2.803, *p*<0.01; GluA2: 260.3±14.2%; *n*=12, *Z*=2. 803, *p*<0.01) and TRPV4 (TRPV4: 293.2±25.4%; GluA1: 209.9±14.6%, *n*=11, *Z*=-2.401, *p*<0.05; GluA2: 276.2±24.3%, *n*=11, *Z*=-1.607, *p*=0.29) compared to GluA1 and GluA2. Asterisks indicate *p*<0.05 (Wilcoxon tests).

**Figure S4.**
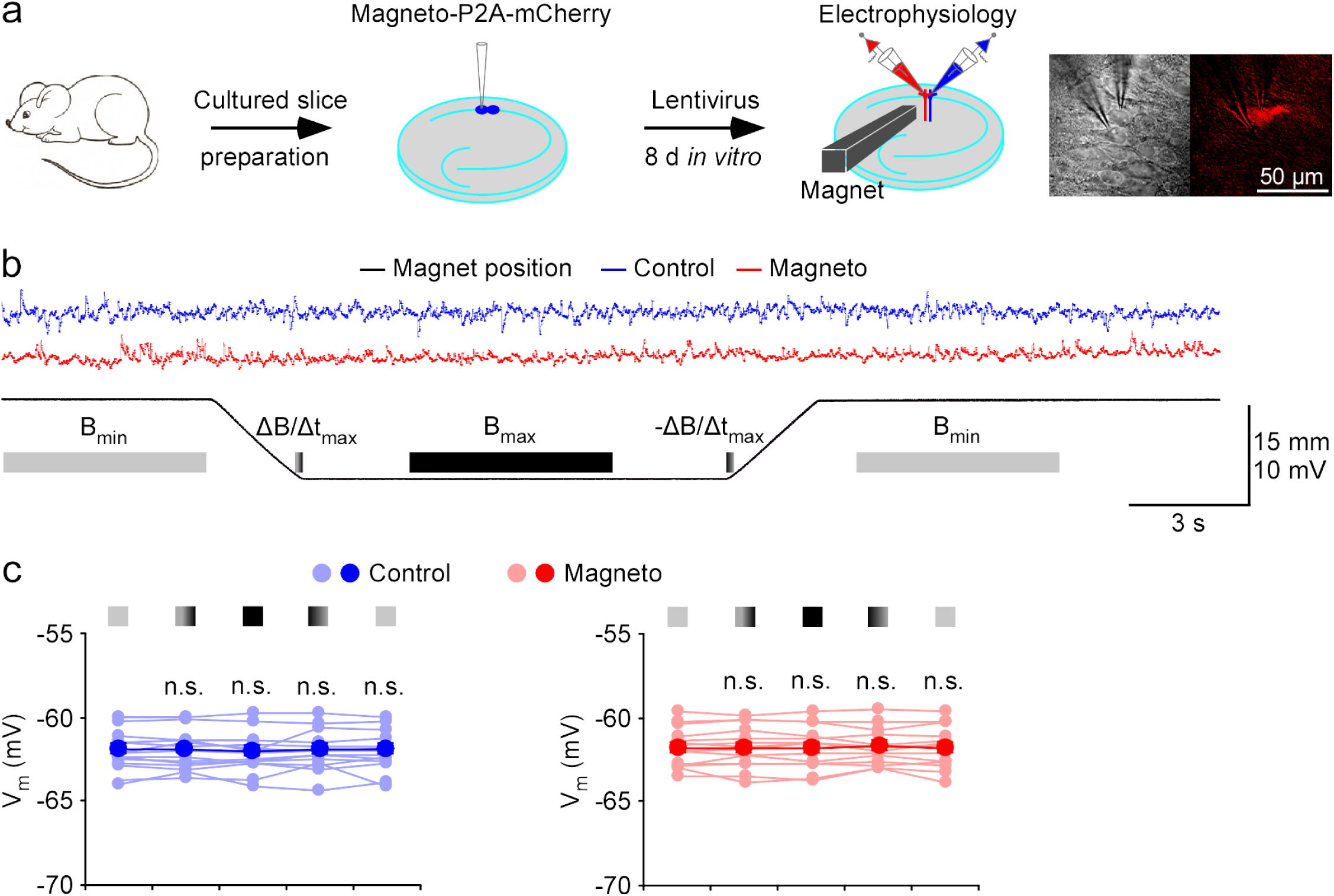
No magnetic effect in rat CA1 neurons expressing lentivirus-delivered Magneto2.0. (a) Schematic drawing outlines the design of *in vitro* lentiviral expression, magnetic stimulation and electrophysiological recordings in cultured rat hippocampal slices. The right images show simultaneous whole-cell recordings from a pair of control non-expressing and Magneto-P2A-mCherry expressing CA1 pyramidal neurons under transmitted light (left) and fluorescence microscopy with RFP filter (right). (b) Recordings of membrane potentials of the pair of control non-expressing and Magneto-P2A-mCherry expressing CA1 pyramidal neurons before, during and after magnetic stimuli delivered with a K&J N42 1/16” permanent block magnet mounted on a micromanipulator. (c) Values of membrane potentials of control non-expressing (Initial B_min_: −62.0±0.3 mV; ΔB/Δt_max_: −61.9±0.3 mV, *Z*=-0.220, *p*=0.83; B_max_: −62.0±0.3 mV, *Z*=-0.471, *p*=0.64; -ΔB/Δt_max_: −61.9±0.3 mV, *Z*=0.282, *p*=0.78; Ending B_min_: −61.9±0.3 mV, *Z*=1.287, *p*=0.20; Wilcoxon tests) and Magneto-P2A-mCherry (Initial B_min_: −61.8±0.3 mV; ΔB/Δt_max_: −61.8±0.3 mV, *Z*=0.282, *p*=0.78; B_max_: −61.8±0.3 mV, *Z*=0.659, *p*=0.51; -ΔB/Δt_max_: −61.7±0.3 mV, *Z*=-0.220, *p*=0.83; Ending B_min_: −61.8±0.3 mV, *Z*=0.345, *p*=0.73; Wilcoxon tests) expressing CA1 pyramidal neurons when the permanent magnet was away from (light), approaching to (light-dark transient color), close to (dark), retracting from (dark-light transient color), and away from (light) recorded neurons (*n*=14 from 6 animals). Note dark lines/dotes indicating averages and shaded lines/dotes indicating experimental data. Note also no difference in membrane potential in control non-expressing and Magneto-P2A-mCherry expressing CA1 pyramidal neurons in all the experimental stages (*p*>0.05; Wilcoxon tests).

**Figure S5.**
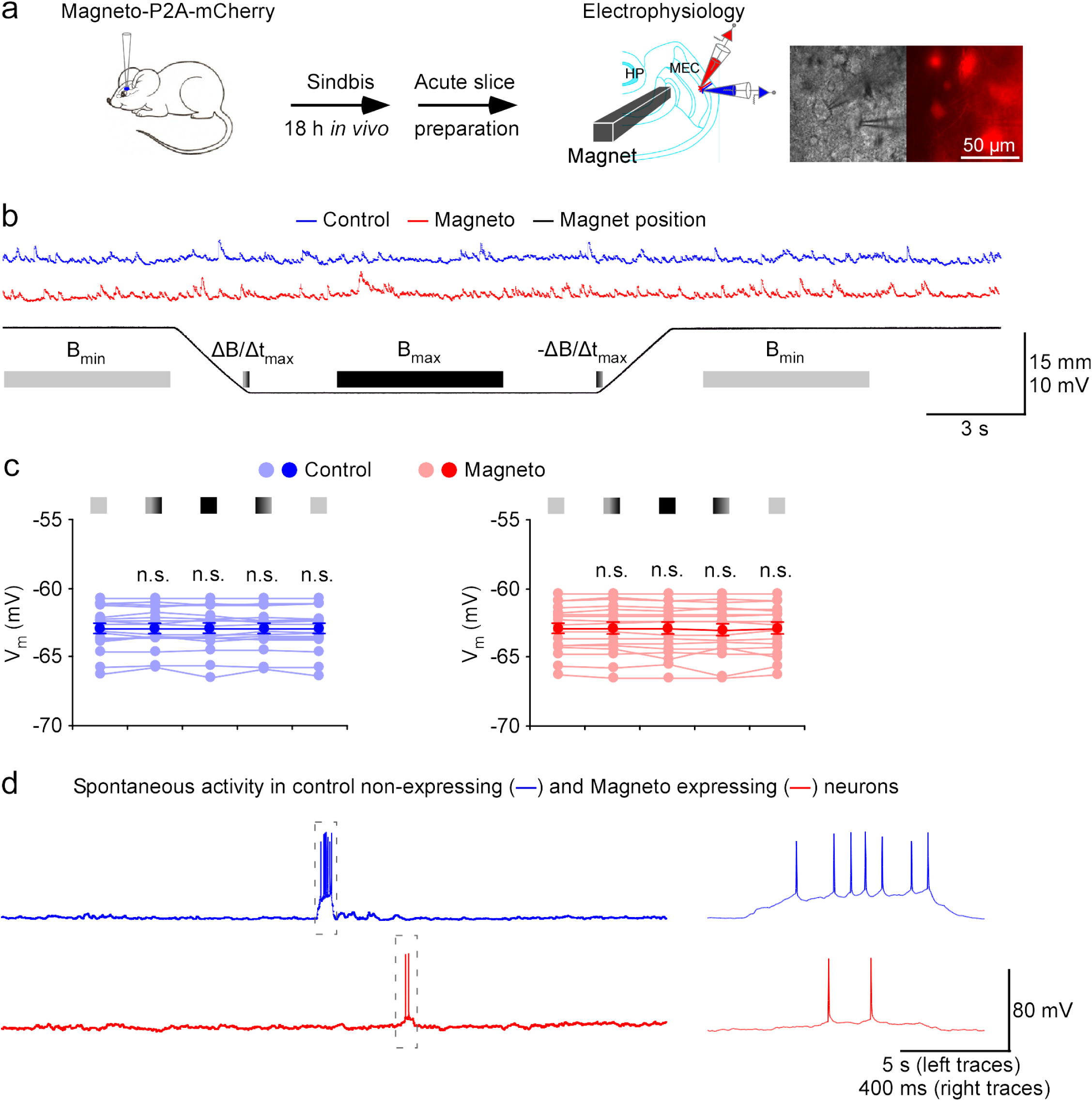
No magnetic effect in mouse MEC L2/3 neurons expressing Sindbis-delivered Magneto2.0. (a) Schematic drawing outlines the design of *in vivo* Sindbis viral expression, *ex vivo* magnetic stimulation and electrophysiological recordings in acutely prepared mouse MEC slices. The right images show simultaneous whole-cell recordings from a pair of control non-expressing and Magneto-P2A-mCherry expressing MEC layer 2/3 pyramidal neurons under transmitted light (left) and fluorescence microscopy with RFP middle filter (right). (b) Recordings of membrane potentials of the pair of control non-expressing and Magneto expressing MEC layer 2/3 neurons before, during and after magnetic stimuli delivered with a K&J N42 1/16” permanent block magnet mounted on a micromanipulator. (c) Values of membrane potentials of control non-expressing (Initial B_min_: −62.8±0.4 mV; ΔB/Δt_max_: −62.8±0.4 mV, *Z*=-0.024, *p*=0.98; B_max_: −62.8±0.4 mV, *Z*=-0.828, *p*=0.41; -ΔB/Δt_max_: −62.8±0.4 mV, *Z*=-0.260, *p*=0.80; Ending B_min_: −62.8±0.4 mV, *Z*=-0.923, *p*=0.36) and Magneto-P2A-mCherry (Initial B_min_: −62.9±0.4 mV; ΔB/Δt_max_: −62.9±0.4 mV, *Z*=0.402, *p*=0.69; B_max_: −63.0±0.4 mV, *Z*=-0.166, *p*=0.87; -ΔB/Δt_max_: −63.0±0.5 mV, *Z*=-1.065, *p*=0.29; Ending B_min_: −63.0±0.4 mV, *Z*=0.166, *p*=0.87) expressing MEC L2/3 pyramidal and stellate neurons when the permanent magnet was away from (light), approaching to (light-dark transient color), close to (dark), retracting from (dark-light transient color), and away from (light) recorded neurons (*n*=17 from 13 animals). Note dark lines/dotes indicating averages and shaded lines/dotes indicating experimental data. Note also no difference in membrane potential in control non-expressing and Magneto-P2A-mCherry expressing MEC L2/3 neurons in all the experimental stages (*p*>0.05; Wilcoxon tests). (d) Recordings of spontaneous events in the pair of control non-expressing and Magneto-P2A-mCherry expressing MEC layer 2/3 neurons. Note that the spontaneous suprathreshold events in the gray dash line boxes are shown again in an expanded time scale in the right.

**Figure S6.**
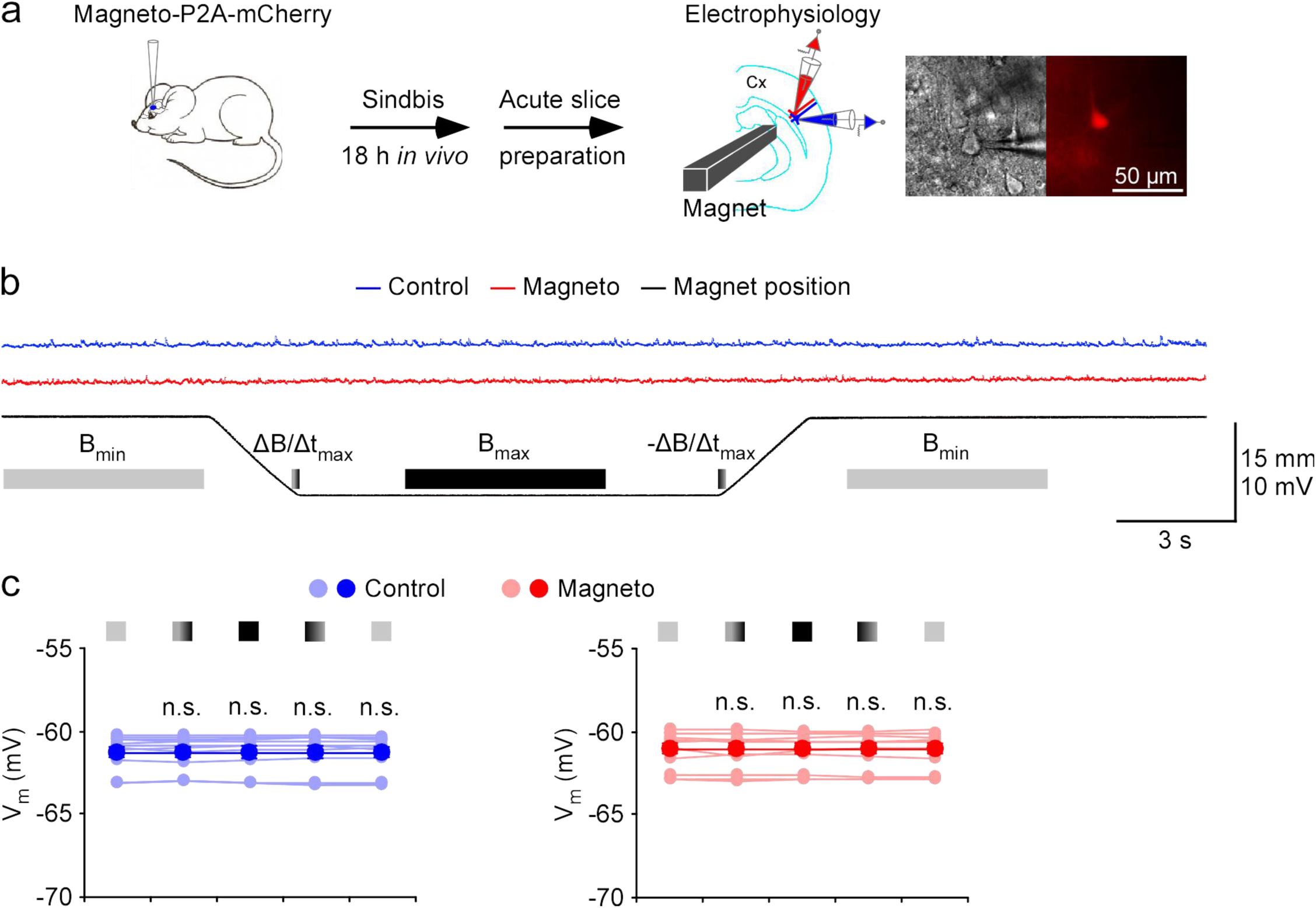
No magnetic effect in mouse barrel cortical L5 neurons expressing Magneto2.0. (a) Schematic drawing outlines the design of *in vivo* Sindbis viral expression, *ex vivo* magnetic stimulation and electrophysiological recordings in acutely prepared mouse cortical slices. The right images show simultaneous whole-cell recordings from a pair of control non-expressing and Magneto-P2A-mCherry expressing barrel cortical L5 pyramidal neurons under transmitted light (left) and fluorescence microscopy with RFP middle filter (right). Cx: cerebral cortex. (b) Recordings of membrane potentials of the pair of control non-expressing and Magneto-P2A-mCherry expressing barrel cortical L5 pyramidal neurons before, during and after magnetic stimuli delivered with a K&J N42 1/16” permanent block magnet mounted on a micromanipulator. (c) Values of membrane potentials of control non-expressing (Initial B_min_: −61.2±0.3 mV; ΔB/Δt_max_: −61.2±0.3 mV, *Z*=0.235, *p*=0.81; B_max_: −61.2±0.3 mV, *Z*=0.000, *p*=1.00; -ΔB/Δt_max_: −61.2±0.3 mV, *Z*=0.628, *p*=0.53; Ending B_min_: −61.2±0.3 mV, *Z*=0.392, *p*=0.70; Wilcoxon tests) and Magneto-P2A-mCherry (Initial B_min_: −61.1±0.3 mV; ΔB/Δt_max_: −61.1±0.3 mV, *Z*=0.000, *p*=1.00; B_max_: −61.1±0.3 mV, *Z*=-0.785, *p*=0.43; -ΔB/Δt_max_: −61.1±0.3 mV, *Z*=-0.235, *p*=0.81; Ending B_min_: −61.1±0.3 mV, *Z*=0.000, *p*=1.00; Wilcoxon tests) expressing barrel cortical pyramidal neurons when the permanent magnet was away from (light), approaching to (light-dark transient color), close to (dark), retracting from (dark-light transient color), and away from (light) recorded neurons (*n*=12 from 6 animals). Note dark lines/dotes indicating averages and shaded lines/dotes indicating experimental data. Note also no difference in membrane potential in control non-expressing and Magneto-P2A-mCherry expressing barrel cortical L5 pyramidal neurons in all the experimental stages (*p*≥0.05; Wilcoxon tests).

**Figure S7.**
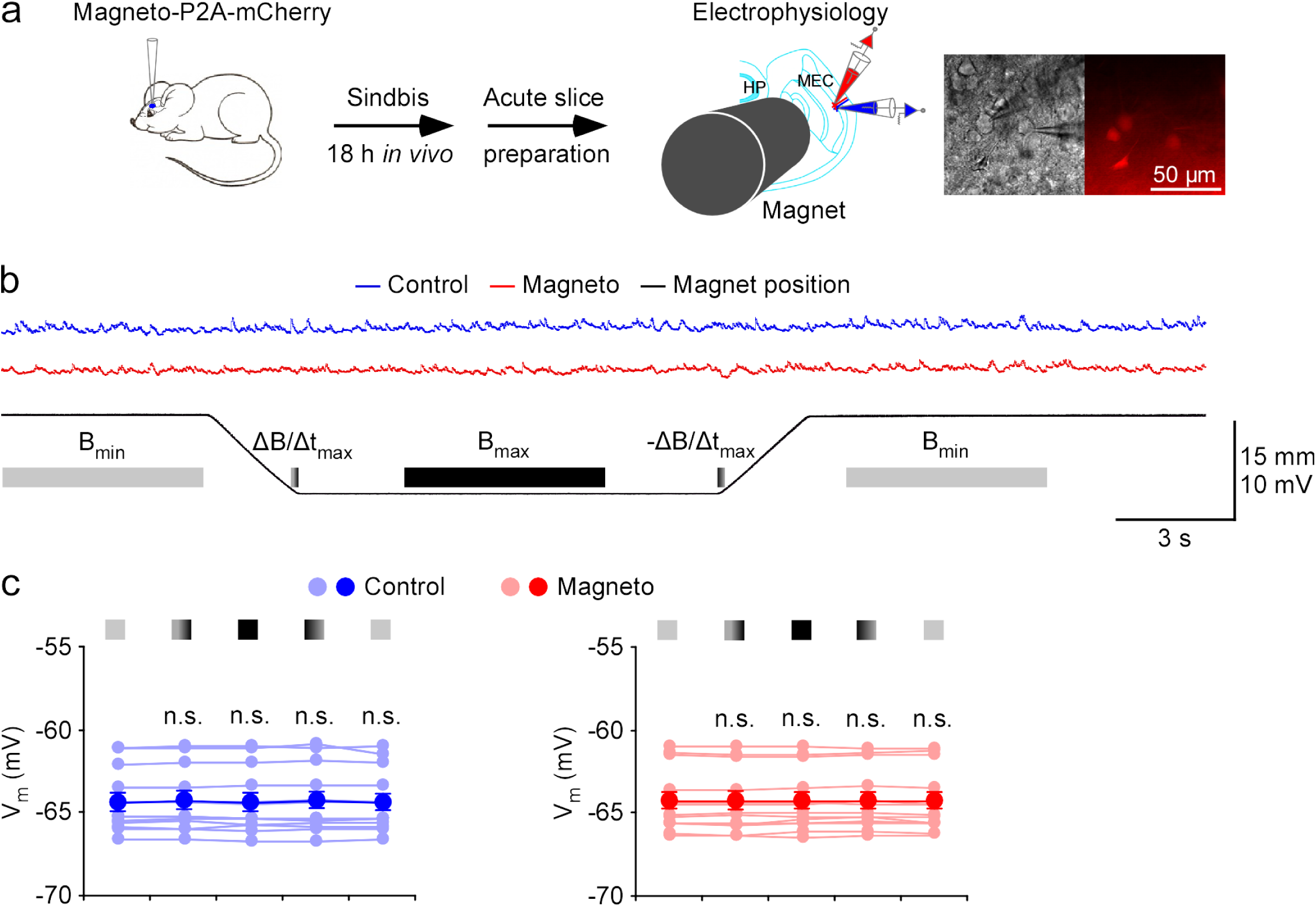
No 3/8” magnet-evoked effect in mouse entorhinal L2/3 neurons expressing Magneto2.0. (a) Schematic drawing outlines the design of *in vivo* Sindbis viral expression, *ex vivo* magnetic stimulation and electrophysiological recordings in acutely prepared mouse MEC slices. The right images show simultaneous whole-cell recordings from a pair of control non-expressing and Magneto-P2A-mCherry expressing MEC L2/3 pyramidal neurons under transmitted light (left) and fluorescence microscopy with RFP middle filter (right). HP: hippocampus; MEC: medial entorhinal cortex. (b) Recordings of membrane potentials of the pair of control non-expressing and Magneto-P2A-mCherry expressing MEC L2/3 pyramidal and stellate neurons before, during and after magnetic stimuli delivered with a 3/8” permanent block magnet mounted on a micromanipulator. (c) Values of membrane potentials of control non-expressing (Initial B_min_: −64.2±0.5 mV; ΔB/Δt_max_: −64.2±0.5 mV, *Z*=0.943, *p*=0.35; B_max_: −64.2±0.5 mV, *Z*=-0.314, *p*=0.75; -ΔB/Δt_max_: −64.2±0.5 mV, *Z*=-1.363, *p*=0.17; Ending B_min_: −64.3±0.5 mV, *Z*=-0.035, *p*=0.97; Wilcoxon tests) and Magneto-P2A-mCherry (Initial B_min_: −64.3±0.5 mV; ΔB/Δt_max_: −64.3±0.5 mV, *Z*=-0.943, *p*=0.35; B_max_: −64.3±0.5 mV, *Z*=0.594, *p*=0.55; -ΔB/Δt_max_: −64.3±0.5 mV, *Z*=1.223, *p*=0.22; Ending B_min_: −64.3±0.5 mV, *Z*=0.524, *p*=0.60; Wilcoxon tests) expressing MEC L2/3 pyramidal and stellate neurons when the permanent magnet was away from (light), approaching to (light-dark transient color), close to (dark), retracting from (dark-light transient color), and away from (light) recorded neurons (*n*=13 from 4 animals). Note dark lines/dotes indicating averages and shaded lines/dotes indicating experimental data. Note also no difference in membrane potential in control non-expressing and Magneto-P2A-mCherry expressing MEC L2/3 neurons in all the experimental stages (*p*>0.05; Wilcoxon tests).

**Figure S8.**
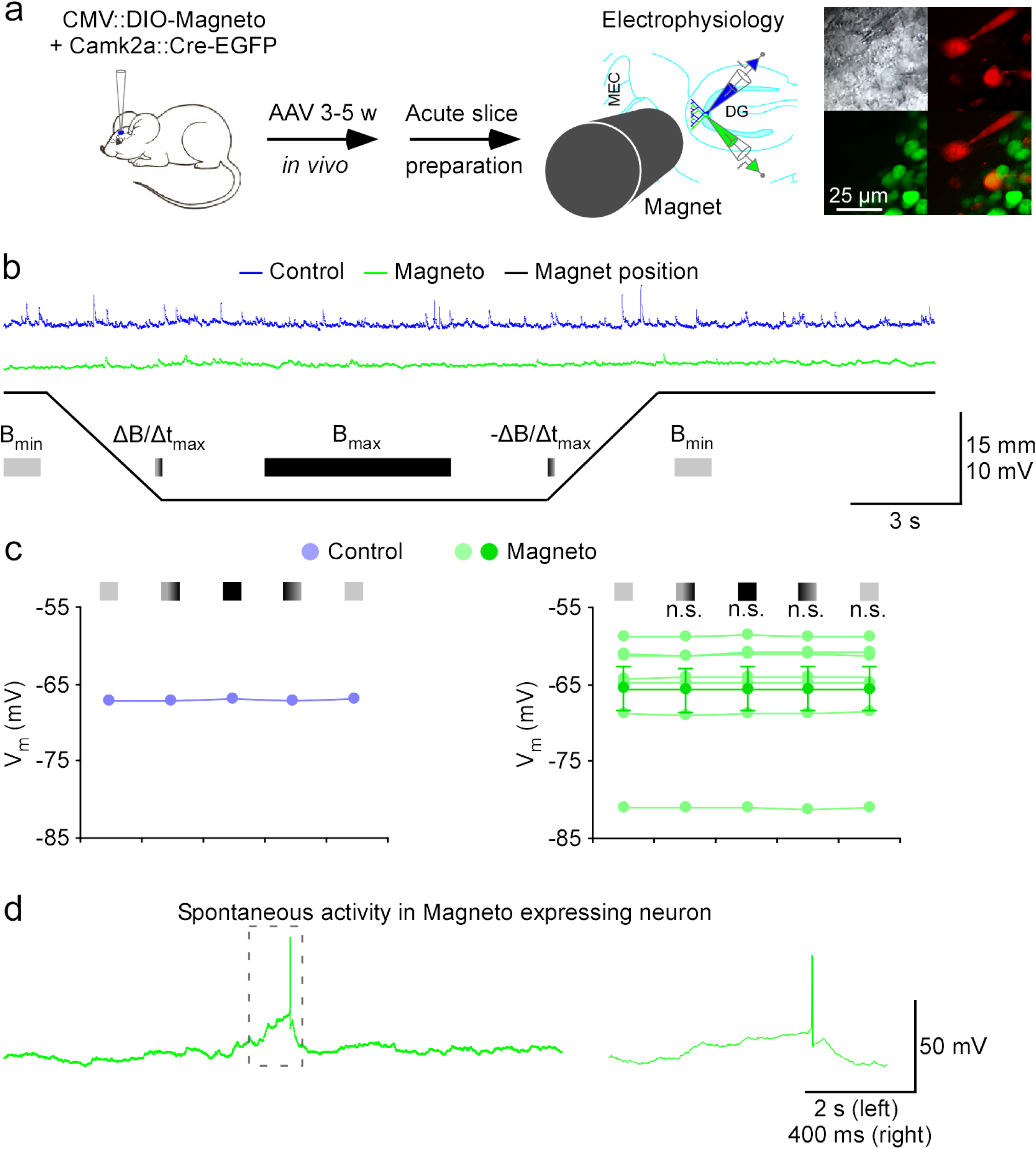
No magnetic effect in mouse hippocampal dentate gyrus neurons expressing Magneto2.0. (a) Schematic drawing outlines the design of *in vivo* AAV viral expression of CMV::DIO-Magneto and Camk2a::Cre-EGFP, *ex vivo* magnetic stimulation and electrophysiological recordings in acutely prepared mouse hippocampal slices. The right images show simultaneous whole-cell recordings from a pair of control non-expressing and DIO-Magneto/Cre-EGFP expressing hippocampal dentate gyrus (**DG**) neurons under transmitted light (upper left) and fluorescence microscopy with RFP (upper right) and GFP (lower left) filters and their overlay (lower right). DG: dentate gyrus; MEC: medial entorhinal cortex. Note that the viral expression and acute slice preparation were carried out by Drs Ronald Gaykema and Manoj Patel. (b) Recordings of membrane potentials of the pair of control non-expressing and DIO-Magneto/Cre-EGFP expressing DG neurons before, during and after magnetic stimuli delivered with a 3/8” permanent block magnet mounted on a micromanipulator. (c) Values of membrane potentials of control non-expressing (Initial B_min_: −66.9 mV; ΔB/Δt_max_: −66.8 mV; B_max_: −66.8 mV; -ΔB/Δt_max_: −66.9 mV; Ending B_min_: −66.8 mV; *n*=1) and DIO-Magneto/Cre-EGFP (Initial B_min_: −65.6±2.8 mV; ΔB/Δt_max_: −65.7±2.8 mV, *Z*=-1.352, *p*=0.18; B_max_: −65.6±2.9 mV, *Z*=1.181, *p*=0.24; -ΔB/Δt_max_: −65.6±2.9 mV, *Z*=0.854, *p*=0.40; Ending B_min_: −65.6±2.8 mV, *Z*=0.854, *p*=0.40; *n*=7) expressing DG neurons when the permanent magnet was away from (light), approaching to (light-dark transient color), close to (dark), retracting from (dark-light transient color), and away from (light) recorded neurons. Note dark lines/dotes indicating averages and shaded lines/dotes indicating experimental data. Note also that in this experiment only one control non-expressing neurons was recorded and analyzed with seven DIO-Magneto/GFP-Cre expressing neurons because the majority of dentate gyrus neurons exhibited green fluorescence (**see insets**). (d) Recordings of spontaneous events in a DIO-Magneto/Cre-EGFP expressing DG neuron. Note that the spontaneous suprathreshold event in the gray dash line box is shown again in an expanded time scale in the right.

**Figure S9.**
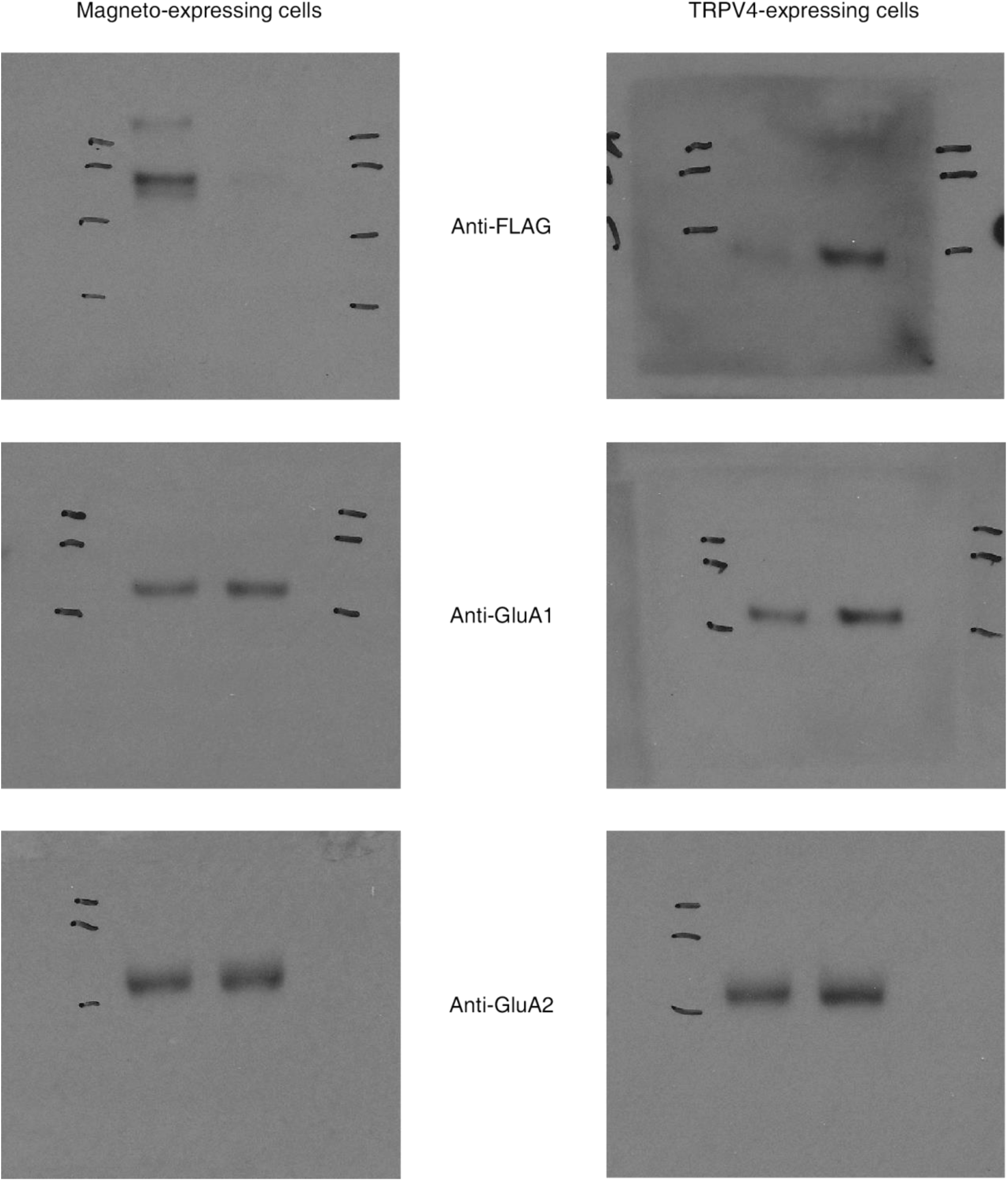
Images of original uncropped Western blots used for preparation of figure 1f-g.

**Figure S10.**
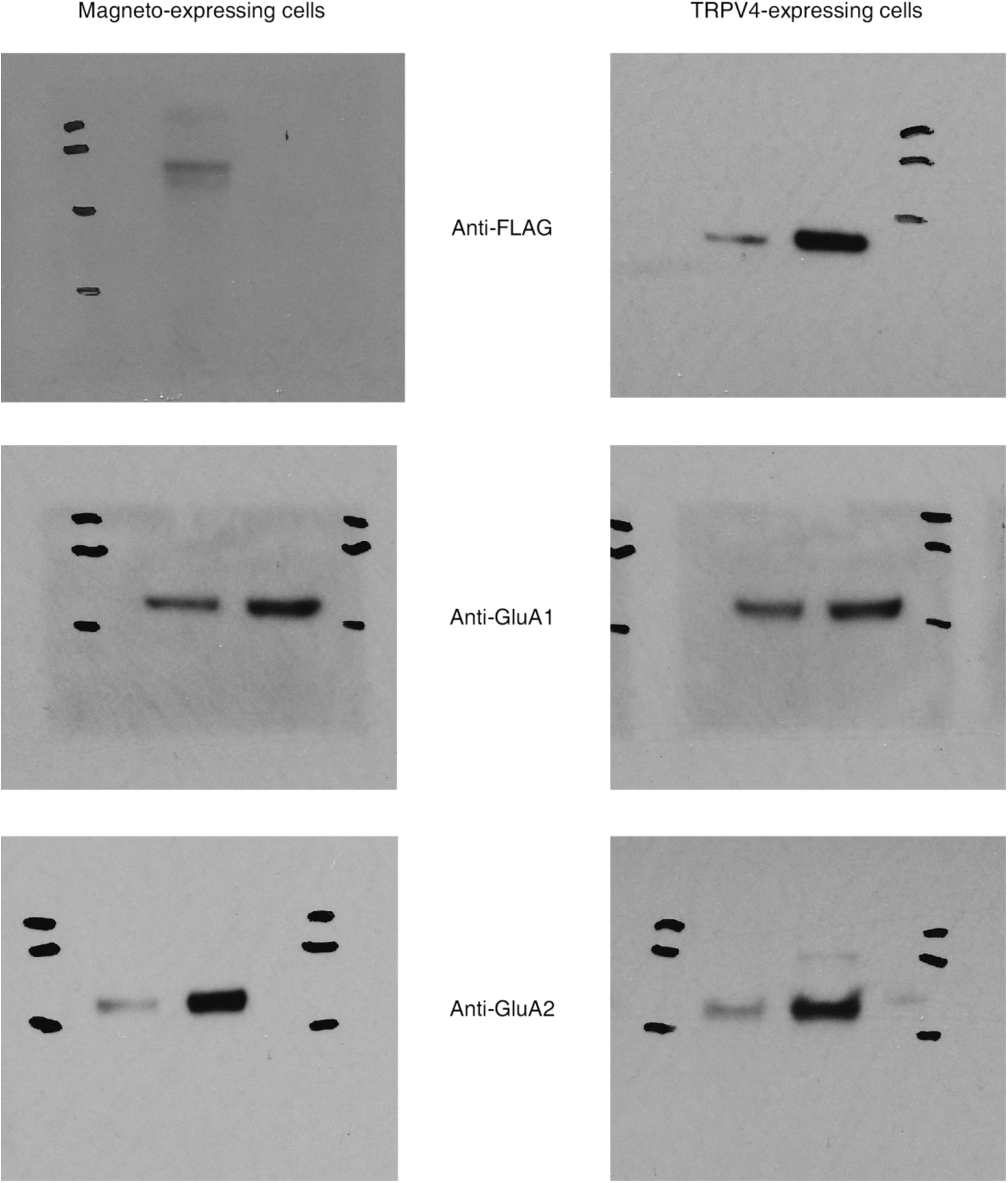
Images of original uncropped Western blots used for preparation of figure S3.

